## Supplementary Material for "SpyMask Enables Combinatorial Assembly of Bispecific Binders"

SpyTag003

RGVPHIVMVDAYKRYK

Masked SpyCatcher003

MSYYHHHHHHDYDIPTTGAMVTTLSGLSGEQGPSGDMTTEEDSATHIKFSKRDEDGRELAGATMELRDSSGKTISTWIS
DGHVKDFYLYPGKYTFVETAAPDGYEVATPIEFTVNEDGQVTVDGEATEGDAGSSGSENLYFQGGGSG**RGVPHIVMVAA
YKRYK****

DoubleCatcher

MSYYHHHHHHDYDIPTTENLYFQGGAMVTTLSGLSGEQGPSGDMTTEEDSATHIKFSKRDEDGRELAGATMELRDSSGK
TISTWISDGHVKDFYLYPGKYTFVETAAPDGYEVATPIEFTVNEDGQVTVDGEATEGDAHTGSGGSGGSGVTTLSGLSG
EQGPSGDMTTEEDSATHIKFSKRDEDGRELAGATMELRDSSGKTISTWISDGHVKDFYLYPGKYTFVETAAPDGYEVAT
PIEFTVNEDGQVTVDGEATEGDAGSSGSENLYFQGGGSG**RGVPHIVMVAAYKRYK***

DoubleCatcher H-Lock

MSYYHHHHHHDYDIPTTENLYFQGGAMVTTLSGLSGEQGPSGDMTTEEDSATHIKFSKRDEDGRELAGATMELRDSSGK
TISTWISDGHVKDFYLYPGKYTFVETAAPDGYEVATPIEFTVNEDGQVTVDGEATEGDAHTGSPANLKALEAQKQKEQR
QAAEELANAKKLKEQLEKGSVTTLSGLSGEQGPSGDMTTEEDSATHIKFSKRDEDGRELAGATMELRDSSGKTISTWIS
DGHVKDFYLYPGKYTFVETAAPDGYEVATPIEFTVNEDGQVTVDGEATEGDAGSSGSENLYFQGGGSG**RGVPHIVMVAA
YKRYK***

DoubleCatcher α-Lock

MSYYHHHHHHDYDIPTTENLYFQGGAMVTTLSGLSGEQGPSGDMTTEEDSATHIKFSKRDEDGRELAGATMELRDSCGK
TISTWISDGHVKDFYLYPGKYTFVETAAPDGYEVATPIEFTVNEDGQVTVDGEATEGDAHTGSGGSGGSGVTTLSGLSG
EQGPSGDMTTEEDSATHIKFSKRDEDGRELAGATMELRDSCGKTISTWISDGHVKDFYLYPGKYTFVETAAPDGYEVAT
PIEFTVNEDGQVTVDGEATEGDAGSSGSENLYFQGGGSG**RGVPHIVMVAAYKRYK***

DoubleCatcher β-Lock

MSYYHHHHHHDYDIPTTENLYFQGGAMVTTCSGLSGEQGPSGDMTTEEDSATHIKFSKRDEDGRELAGATMELRDSSGK
TISTWISDGHVKDFYLYPGKYTFVETAAPDGYEVATPIEFTVNEDGQVTVDGEATEGDAHTGSGGSGGSGVTTLSGLSG
EQGPCGDMTTEEDSATHIKFSKRDEDGRELAGATMELRDSSGKTISTWISDGHVKDFYLYPGKYTFVETAAPDGYEVAT
PIEFTVNEDGQVTVDGEATEGDAGSSGSENLYFQGGGSG**RGVPHIVMVAAYKRYK***

DoubleCatcher γ-Lock

MSYYHHHHHHDYDIPTTENLYFQGGAMVTTLSGLSGEQCPSGDMTTEEDSATHIKFSKRDEDGRELAGATMELRDSSGK
TISTWISDGHVKDFYLYPGKYTFVETAAPDGYEVATPIEFTVNEDGQVTVDGEATEGDAHTGSGGSGGSGVTTLSGLSG
EQGPSGDMTTEEDSATHIKFSKRDEDGRELAGATMELRDSSCKTISTWISDGHVKDFYLYPGKYTFVETAAPDGYEVAT
PIEFTVNEDGQVTVDGEATEGDAGSSGSENLYFQGGGSG**RGVPHIVMVAAYKRYK***

DoubleCatcher δ-Lock

MSYYHHHHHHDYDIPTTENLYFQGGAMVTTLSGLSGEQGPSGDMTTEEDSATHIKFSKRDEDGRELAGATMELRDSSGK
TISTWISDGHVKDFYLCPGKYTFVETAAPDGYEVATPIEFTVNEDGQVTVDGEATEGDAHTGSGGSGGSGVTTLSGLSG
EQGPSGDMTTEEDSATHIKFSKRDEDGRELAGATMELRDSSGKTISTWISDGHVKDFYLCPGKYTFVETAAPDGYEVAT
PIEFTVNEDGQVTVDGEATEGDAGSSGSENLYFQGGGSG**RGVPHIVMVAAYKRYK***

DoubleCatcher ε-Lock

MSYYHHHHHHDYDIPTTENLYFQGGAMVTTLSGLSGEQGPSGDMTTEEDSATHIKFSKRDEDGRELAGATMELRDSSGC
TISTWISDGHVKDFYLYPGKYTFVETAAPDGYEVATPIEFTVNEDGQVTVDGEATEGDAHTGSGGSGGSGVTTLSGLSG
EQGPSGDMTTEEDSATHIKFSKRDEDGRELAGATMELRDSSGKTISTWISDGCVKDFYLYPGKYTFVETAAPDGYEVAT
PIEFTVNEDGQVTVDGEATEGDAGSSGSENLYFQGGGSG**RGVPHIVMVAAYKRYK***

***Supplementary Figure 1.* Amino acid sequences of SpyTag003, Masked SpyCatcher003, and DoubleCatcher variants.** The $His_6$-tag is shown with gray shading, TEV protease cleavage site with green shading, GSG spacer with cyan shading, cysteines in red, SpyTag003DA mask in bold, and the stop codon with *.

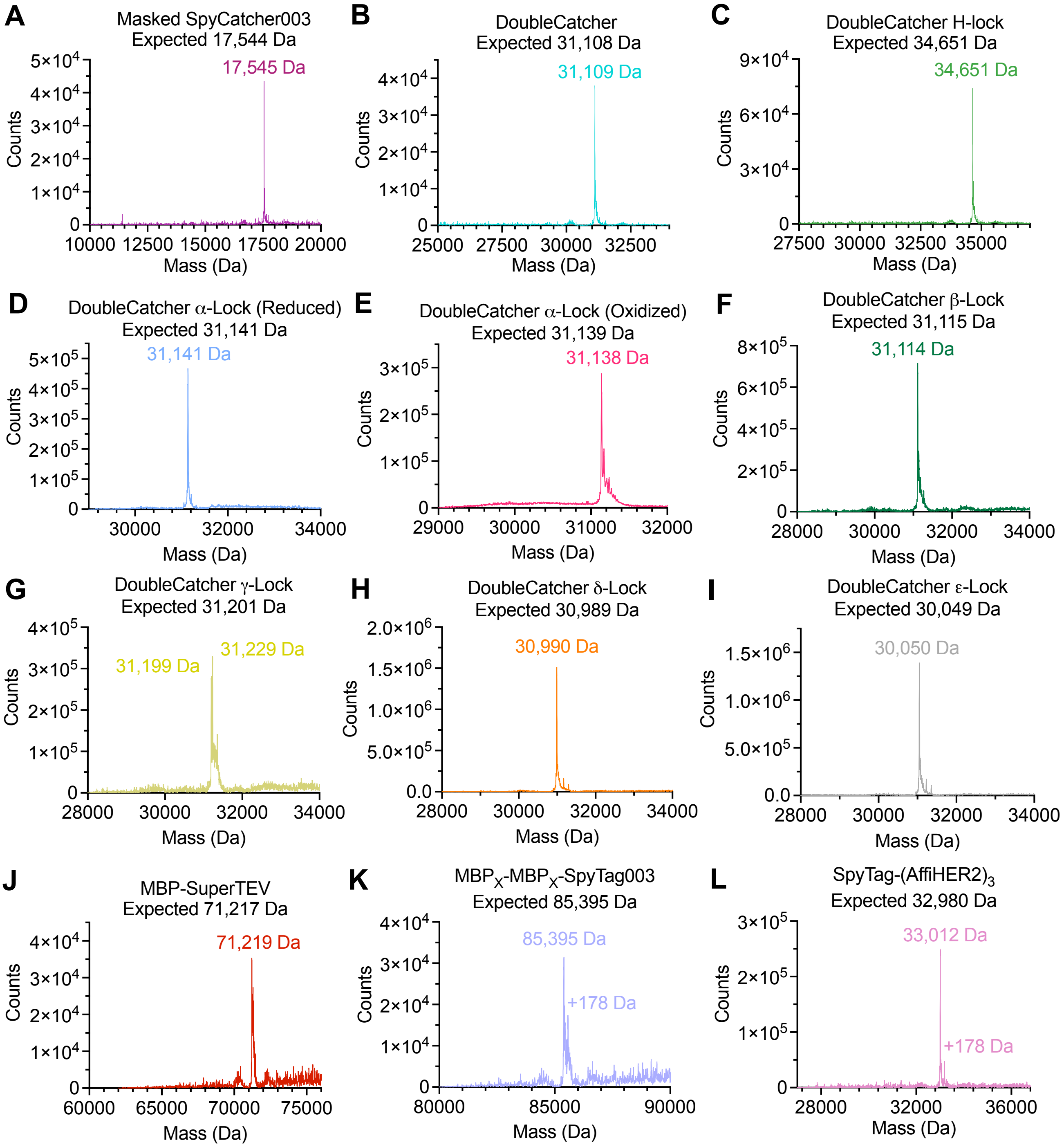


***Supplementary Figure 2.*** **Mass spectrometry of building blocks.** RapidFire Electrospray Ionization Mass Spectrometry on Masked SpyCatcher, DoubleCatcher variants and each Tagged binder. The observed mass is indicated above the main peak. The expected mass was calculated from ExPASy ProtParam, based on disulfide bonds being formed. The minor +178 Da peak relates to gluconoylation, which is a common post-translational modification for proteins overexpressed in *E. coli* BL21 (Geoghegan et al., 1999).

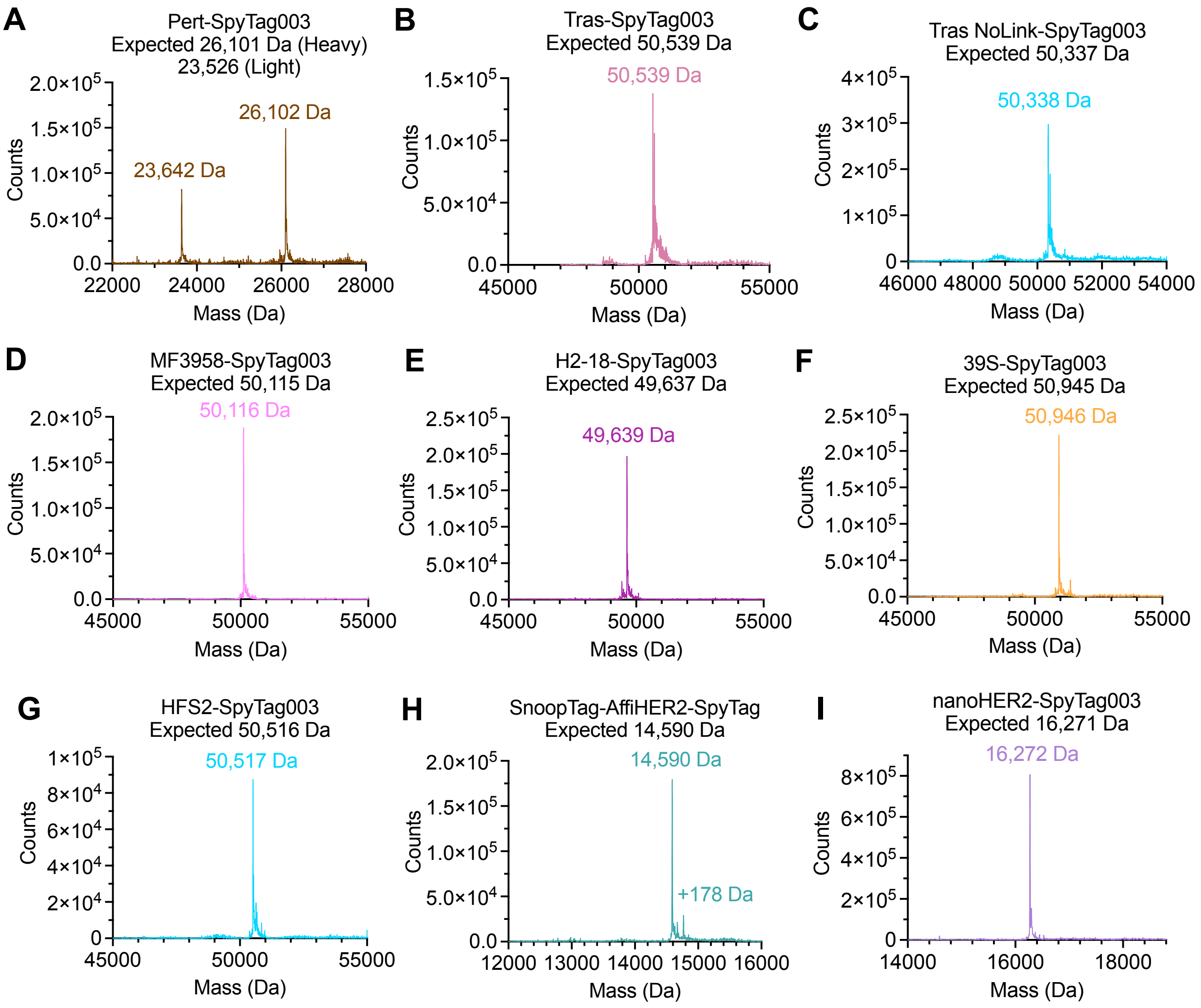


***Supplementary Figure 3.*** **Mass spectrometry of anti-HER2 binders.** RapidFire Electrospray Ionization Mass Spectrometry of SpyTag-linked anti-HER2 Fabs, an affibody, and a nanobody, as used to assemble the matrix of bispecifics. The observed mass is indicated above the main peak. Since the hinge region is omitted from the Pert-SpyTag003 heavy chain sequence, there is no interchain disulfide bond in the Pert Fab. The expected mass was calculated from ExPASy ProtParam. The minor +178 Da peak relates to gluconoylation.

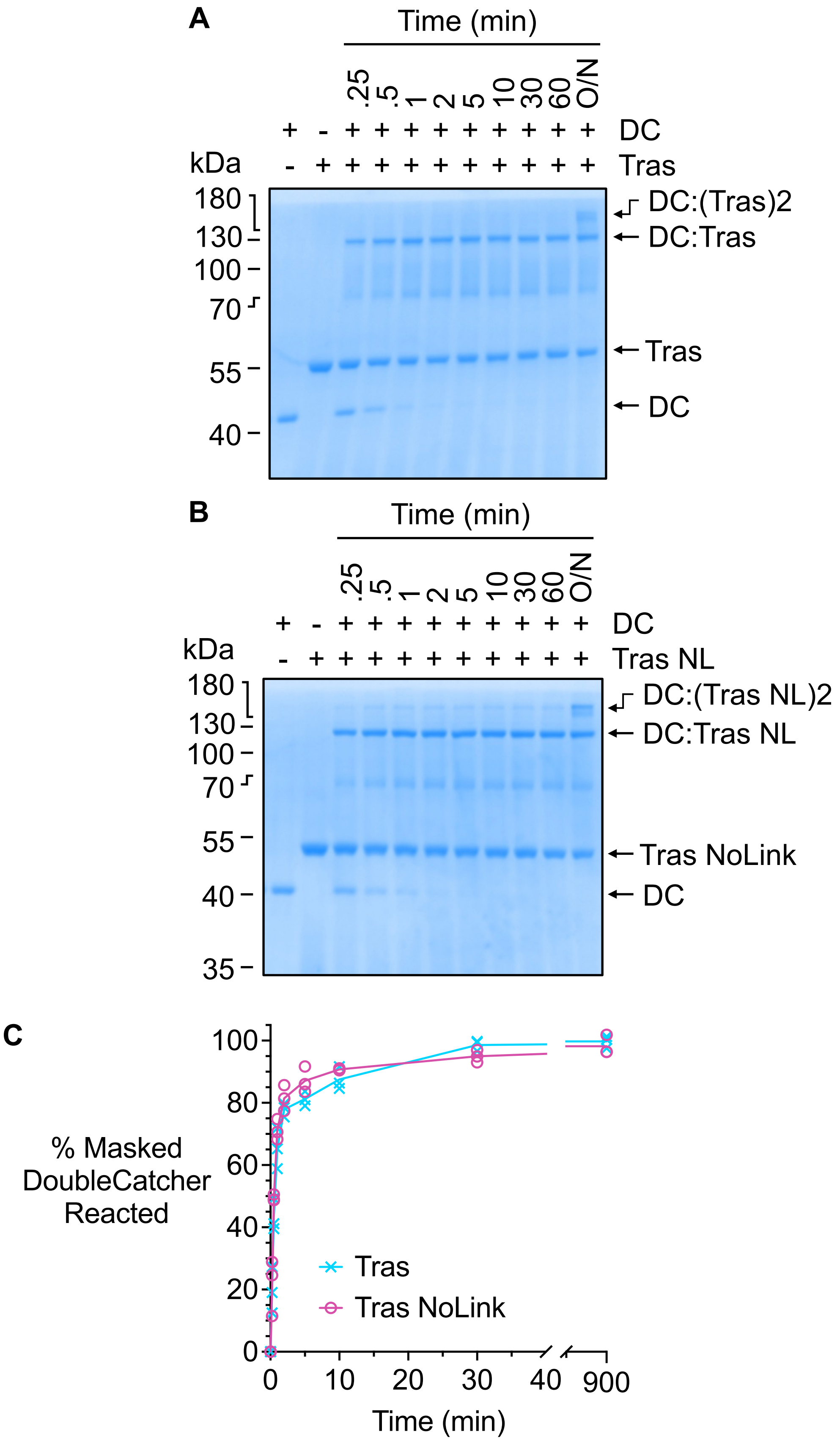


***Supplementary Figure 4.*** **Removing the linker between Tras Fab and SpyTag003 does not affect reactivity with DoubleCatcher.** The tripeptide linker connecting Tras Fab and SpyTag003 was removed to generate Tras NoLink (Tras NL), to reduce the flexibility between the DoubleCatcher and the binder. (**A**) Time-course of reaction between 2.5 μM DoubleCatcher and 5 μM Tras in PBS pH 7.4 at 37 °C for the indicated time, analyzed by SDS-PAGE/Commassie. (**B**) As in (A) with Tras NL. (**C**) Quantification of reactivity of Tras or Tras NoLink for DoubleCatcher. Each datapoint is shown (n = 3), with the line connecting the mean.
